# A DNA translocase operates by cycling between planar and lock-washer structures

**DOI:** 10.1101/2020.05.22.101154

**Authors:** Juan P. Castillo, Alexander Tong, Sara Tafoya, Paul J. Jardine, Carlos Bustamante

## Abstract

Ring ATPases that translocate disordered polymers possess lock-washer architectures that they impose on their substrates during transport via a *hand-over-hand* mechanism. Here, we investigate the operation of ring motors that transport substrates possessing a preexisting helical structure, such as the bacteriophage ϕ29 dsDNA packaging motor. During each cycle, this pentameric motor tracks one helix strand (the ‘tracking strand’), and alternates between two segregated phases: a *dwell* in which it exchanges ADP for ATP and a *burst* in which it packages a full turn of DNA in four steps. We challenge this motor with DNA-RNA hybrids and dsRNA substrates and find that it adapts the size of its burst to the corresponding shorter helical pitches by keeping three of its power strokes invariant while shortening the fourth. Intermittently, the motor loses grip when the tracking strand is RNA, indicating that it makes load-bearing contacts with the substrate that are optimal with dsDNA. The motor possesses weaker grip when ADP-bound at the end of the burst. To rationalize all these observations, we propose a *helical inchworm* translocation mechanism in which the motor increasingly adopts a lock-washer structure during the ATP loading dwell and successively regains its planar form with each power stroke during the burst.

## Introduction

Ring-shaped ATPases of the ASCE division convert the energy of nucleotide hydrolysis into mechanical work to drive a variety of processes inside living cells, from electro-chemical gradient buildup, to protein degradation, and DNA translocation, among others[1]. Various aspects of the operation of ring-shaped motors such as force generation, substrate engagement, mechano-chemical conversion, and inter-subunit coordination have been described[2-9]. In addition, recent high-resolution structures of ring-shaped ATPases (particularly helicases and polypeptide translocases) have revealed lock-washer architectures[10-12]. Based on these structures, a *hand-over-hand* mechanism of translocation has been proposed, in which the crack in the ring precesses along the subunits, and the motor imposes its helical structure on the otherwise disordered substrate. The question then arises as to what is the mechanism of operation of ring ATPases that adopt lock-washer conformations but translocate structured substrates, such as dsDNA?

To tackle this question, we investigate how the structure and symmetry properties of the substrate determine the stepping mechanism of the DNA packaging motor of bacteriophage ϕ29, which has recently been shown to adopt a lock-washer structure in which the individual subunits contact the phosphates of the DNA backbone (Marc Morais, personal communication*). This homopentameric motor displays a mechano-chemical cycle of bursts—during which 10 bp of double-stranded DNA (dsDNA) are internalized in the viral capsid—segregated from dwells—wherein the motor subunits exchange ADP for ATP in a sequential fashion around the ring[3]. Moreover, each burst is made of four 2.5 bp steps (0.85 nm), indicating that one of the five subunits is *special* in that it does not perform a mechanical but a regulatory function[5, 7]. This task requires the special subunit to contact a pair of phosphates in the strand that is being packaged in the 5’ to 3’ direction (the *tracking strand*) every pitch [4]. The segregation between phases of nucleotide exchange and hydrolysis-coupled translocation is incompatible with the hand-over-hand mechanism, where nucleotide exchange and hydrolysis are interlaced. Hence, different mechanisms of translocation have been proposed for ϕ29; however, definitive evidence for any of them is missing[3, 13].

To gain insight into this mechanism, we monitored the operation of the ϕ29 packaging motor on double-stranded RNA and DNA/RNA hybrid substrates, whose helical periods are shorter than canonical B-form DNA. Significantly, we find that the motor adapts its burst size to match the new substrate’s periodicity. However, we find that three of the individual steps within the modified burst retain their original size—indicating that these are fixed and determined only by the conformational changes of the subunits concomitant with ATP hydrolysis—whereas the fourth is reduced to match the pitch of the double helix. Finally, although the motor is highly processive when the tracking strand is DNA, it displays burst-sized slipping when it is RNA. Based on these observations, we propose a model for the packaging mechanism in which the ring cracks open, progressively adopting a lock-washer conformation during the ATP loading dwell, and successively returns to a planar structure upon sequential hydrolysis during the translocation burst.

### The motor can translocate A-form substrates

The natural substrate of the ϕ29 packaging motor is dsDNA, which adopts the canonical B-form under physiological conditions. Double-stranded RNA (dsRNA) and DNA/RNA hybrids, in contrast, adopt exclusively the A-form, displaying shorter helical pitches while the main features of their solvent-exposed phosphodiester backbones and base-pair stacking cores remain essentially the same[14-17] (Fig. 1a, Table 1). We challenged the ϕ29 packaging motor with these alternative structures and monitored its translocation at the single molecule level using high-resolution optical tweezers. We designate the DNA/RNA hybrids either as DNA tracking strand (DTS) hybrid or RNA tracking strand (RTS) hybrid, depending on the chemical identity of the strand being packaged in the 5’ to 3’ direction (Fig. 1a).

**Table 1.**
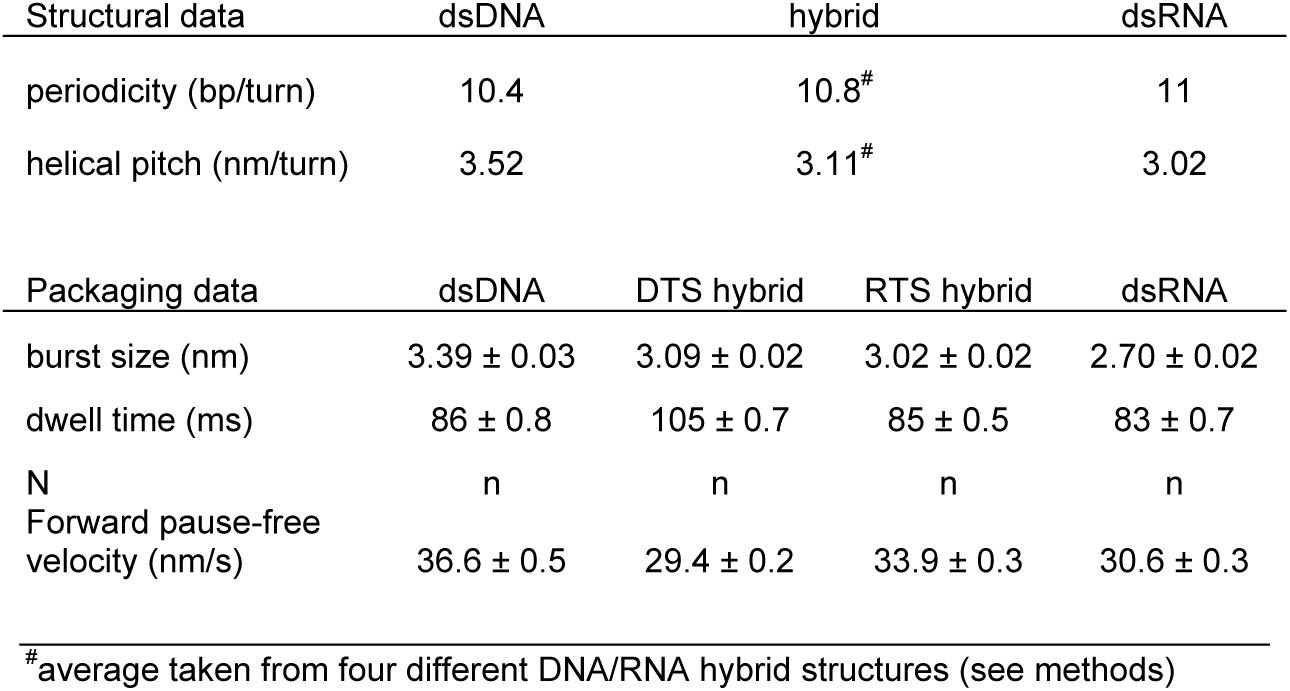
Structural and packaging features of the four different substrates

**Fig. 1.**
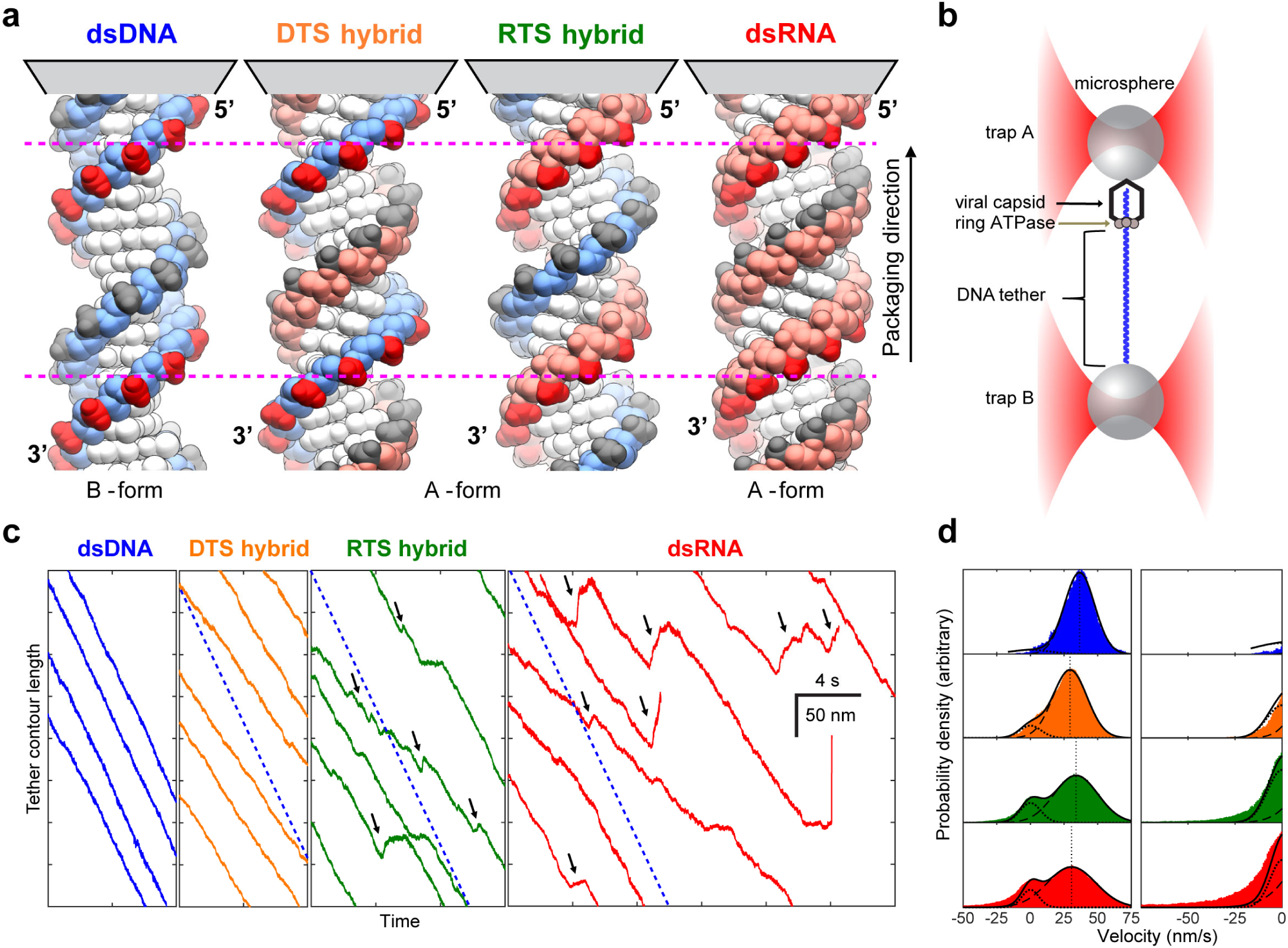
Packaging of alternative substrates. **a**, The structure of the four substrates is shown, with DNA and RNA strands in light blue and light red, respectively. The strand being packaged from 5’ to 3’ is designated the tracking strand, and its phosphates are colored dark red. Note the differing orientation of the phosphates for DNA and RNA strands, and the differing helical pitches (Table 1, red dotted lines span one pitch of dsDNA). **b**, Single packaging complexes are held between two microspheres using a dual trap optical tweezer while the tether length is monitored over time. **c**, Representative packaging trajectories for the four substrates are shown at 250 Hz. Dashed blue lines represent the velocity of packaging on dsDNA substrate. Reverse translocation dynamics are observed on the RTS hybrid and dsRNA (black arrows). **d**, Velocity distributions for the different substrates are fit to the sum of two gaussians (dark line), one with zero mean (dotted line) and one with positive mean (dashed line) that represent pausing and packaging, respectively. The right panel details negative velocities, which correspond to reverse translocation dynamics. Note the increased probability of these events on dsRNA compared to the RTS hybrid. For all panels, [ATP] is 0.25 mM and opposing force is 7-12 pN.

Individual packaging complexes were tethered between two microspheres in a dual trap optical tweezers instrument[3] (Fig. 1b) and the tether’s end-to-end distance was monitored over time while the motor packaged one of the three A-form substrates in the presence of ATP. Fig. 1c depicts the packaging trajectories obtained over these alternative substrates compared with those over dsDNA at saturating [ATP] (0.25 mM ∼ 8 x *K*_*m*_) and 7-12 pN of opposing force[2]. The motor processively packages all A-form substrates, which rules out a recent model of motor operation which proposes that the translocation mechanism involves a B-form to A-form transition of the substrate induced by dehydration and driven by ATP hydrolysis[13, 18], since dsRNA cannot adopt the B-form.

The motor packages the hybrids and dsRNA slower than dsDNA, with average packaging velocities of 36.6 ± 0.5 nm/s for dsDNA, 29.4 ± 0.2 nm/s for the DTS hybrid, 33.9 ± 0.3 nm/s for the RTS hybrid, and 30.6 ± 0.3 nm/s for dsRNA (all values mean ± s.e.m.) (Fig. 1c,d, Table 1). Significantly, when packaging the RTS hybrid and dsRNA, the motor displays reverse translocation events during which the tether length increases with a finite slope for a period of time until eventually it resumes packaging (Figure 1c, black arrows). In contrast, when packaging dsDNA and the DTS hybrid, the motor is extremely processive, suggesting that the cause of the reverse translocation events is weaker motor-substrate interactions when the tracking strand is RNA. Fig. 1d displays the velocity distributions obtained for the different substrates, wherein the reverse translocation dynamics manifest as long tails of negative velocities in the distribution.

### The motor burst adapts to the helical pitch

To understand how the motor processes these different substrates, we analyzed packaging trajectories at high resolution. With all three A-form substrates, the motor conserves its dwell-burst operational scheme (Fig. 2a). The dwell times for the RTS hybrid and dsRNA are similar to those of dsDNA (Table 1). Only the DTS hybrid values show a significant difference—we do not have an explanation for this behavior. Moreover, the dwell times are gamma-distributed on all four substrates, just as it was previously described for dsDNA [3, 5], an indication that nucleotide exchange of the motor subunits is essentially unaltered by the A-form structures (Supplementary Fig. S1 and Supplementary Discussion). Importantly, however, we find that the bursts are smaller for all three A-form substrates (Fig. 2a). The pairwise distance distribution analysis of high-resolution packaging trajectories (Fig. 2b) reveals that the burst size for the A-form substrates is reduced to 3.1 ± 0.02 nm, 3.0 ± 0.02 nm and 2.7 ± 0.02 nm for the hybrids and dsRNA, respectively, from 3.4 ± 0.03 nm for dsDNA (all values mean ± s.e.m.). Significantly, these burst sizes closely match the corresponding helical pitches of the substrates (Table 1).

**Fig. 2.**
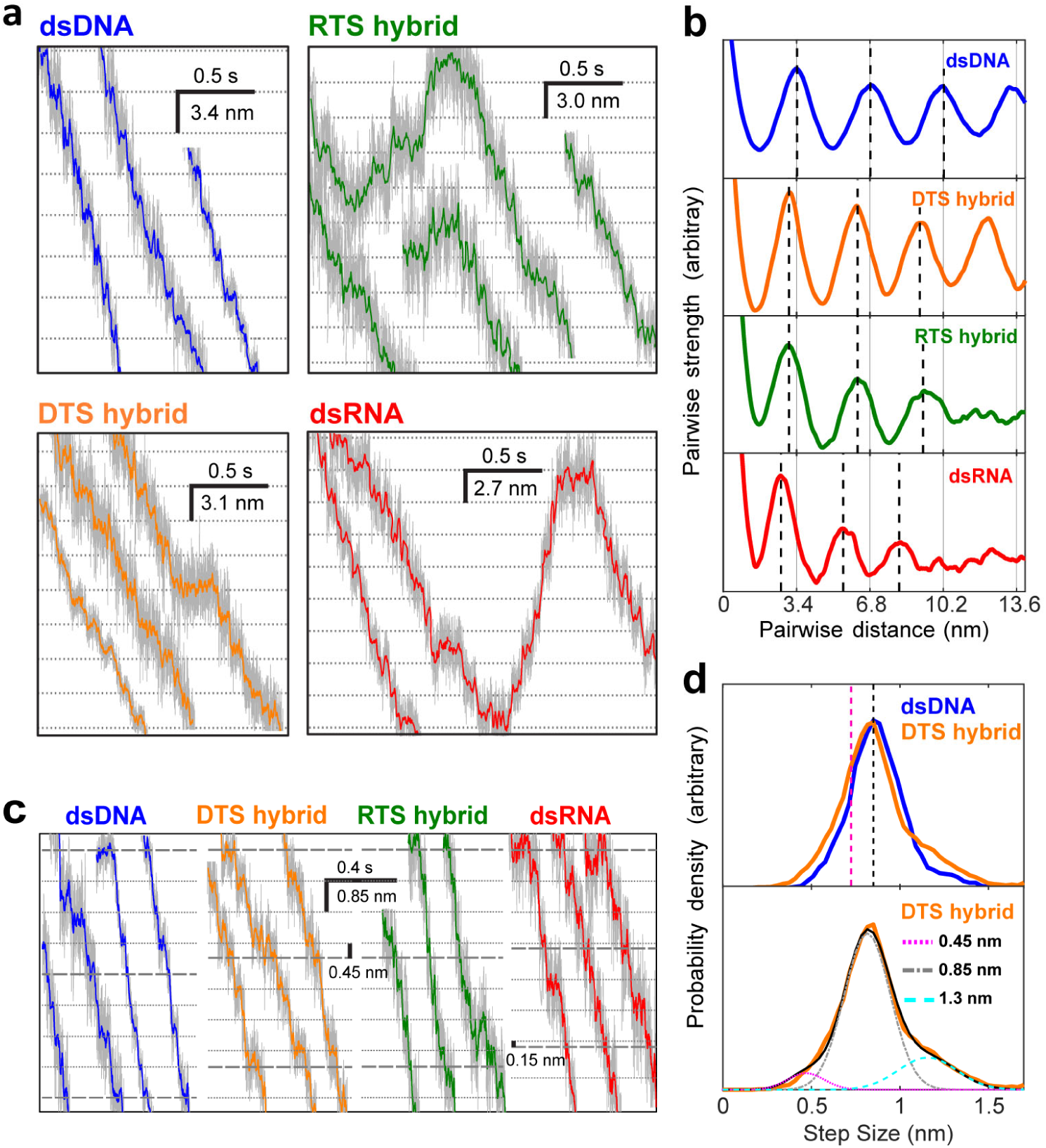
Burst and step sizes for different substrates. **a**, Representative packaging traces at low force (7-12 pN) are shown for the four substrates at 2.5 kHz (grey) and filtered to 100 Hz (colored). On A-form substrates, the dwell-burst coordination of the motor is retained but with a changed burst size. The dashed lines are spaced to match the mean burst size of the substrates. **b**, Pairwise distance analysis of the trajectories confirms a decrease of the burst size for the three A-form substrates. The dashed lines are evenly spaced in multiples of the mean burst size. **c**, Representative traces at high force (35-40 pN) resolves motor steps. For dsDNA, steps of 0.85 nm are observed, however for A-form substrates smaller steps are also seen every fourth step. **d**, Step size distributions from packaging traces against high-force. Top: the step size distribution peaks at 0.85 nm for dsDNA and the DTS hybrid (black dashed line). Magenta dashed line indicates the expected peak if the four elementary steps were instead shortened proportionally with the reduction in helical pitch (0.72 nm). Bottom: DTS hybrid step size distribution accounts for steps of size 0.45 nm, 0.85 nm and their sum (if the two steps were combined during stepfinding) by fitting three gaussians. For all panels, [ATP] is 0.25 mM.

To determine the size of the steps that compose the reduced bursts, we obtained packaging trajectories under 30-35 pN of opposing force. On dsDNA, as previously reported, the burst is made up of four steps of 0.85 nm[3]. On the DTS hybrid, we find that each burst is instead composed of three 0.85 nm steps and one smaller step of 0.45 nm (Fig 2c-d). In addition, we also observe 0.85 nm steps on the RTS hybrid and dsRNA, but slipping frequency and resolution limitations in these substrates prevent the quantitative determination of the smaller steps (Supplementary Discussion). However, since the total size of the burst on the RTS hybrid and dsRNA are 3.0 nm and 2.7 nm, respectively, we determine that the shortened step for the former substrate is 0.45 nm and 0.15 nm for the latter (Fig. 2c).

These results demonstrate that i) the inherent step size of the motor subunits is most likely determined by the conformational change that accompanies their power stroke during phosphate release, and ii) to successfully package a reduced helical pitch substrate, the motor decreases the size of its fourth power stroke during each burst, possibly due to the earlier encounter of the last subunit with the backbone phosphate one pitch away in the helix[4]. Since the cryo-EM lock-washer structure of this motor shows that it spans one pitch of a dsDNA helix (Marc Morais, personal communication*), our results suggest that the motor adapts to the A-form substrates, opening less on them by distorting one of the inter-subunit interfaces.

### Burst-size slipping on RNA tracking strands

Surprisingly, the reverse translocation events observed only on the RTS hybrid and dsRNA occur in increments whose size coincide with the magnitude of the burst on the corresponding substrate (Fig 2a, Supplementary Fig. S2a). Therefore, we refer to them as *burst-sized slips*. The appearance of burst-sized slipping only when with RTS hybrid and dsRNA, suggests that critical interactions between subunits in the motor and the DNA tracking strand confer stable grip during packaging, and that these can be sporadically lost when this strand is RNA. The brief periodic dwells separated by sudden slips presumably reflect the motor weakly gripping the substrate through the same phosphate contacts it makes during a normal packaging dwell[4] (Supplementary Discussion) and releasing it to grip it again one full turn away. We find that the number of burst-sized slips within a reverse translocation event does not increase with limiting [ATP] (10-25 μM), suggesting that nucleotide exchange is halted during the course of an event (see Supplementary Discussion, Supplementary Fig. S2b). Thus, we propose that, to enter a reverse translocation event, the motor must transition into a *packaging-incompetent* state in which it is unable to perform nucleotide exchange, and that interacts weakly with the RNA tracking strand. On the other hand, limiting [ATP] does increase the frequency of events (Supplementary Fig. S2b), consistent with a model where the attainment of the packaging-incompetent state is in kinetic competition with the completion of the ATP loading process; accordingly, faster saturation of the ring with ATP reduces the probability of observing burst-sized slipping (see Supplementary Discussion). Previous studies have shown that this motor and similar viral dsDNA translocases display a decreasing grip for the substrate when subunits in the ring are in the ATP, ADP, and apo forms, respectively[2, 19]. Taken together, the above observations indicate that ATP saturation of the motor confers strong grip for its substrate via the tracking strand. Furthermore, the length of time the motor spends moving backwards during reverse translocation events is 4.1 times longer on a dsRNA substrate than on an RTS hybrid (Supplementary Fig. S2d). This observation indicates that the rate at which the motor regains stable grip is sensitive to the structural details of the tracking strand, presumably the helical pitch, which is shortest for dsRNA, and/or other structural differences between the tracking strands. (Fig. 1a, Table 1).

### A helical inchworm model of translocation

The reduced grip of the motor for substrates harboring an RNA tracking strand of decreasing helical pitch suggests that during translocation the motor adopts a lock-washer conformation in which its subunit interact optimally with the tracking strand of dsDNA. Moreover, the correlation between increased grip and ATP saturation suggests that the interaction between the motor and the tracking strand is progressively established as the subunits in the motor sequentially bind ATP during the dwell. Yet, if the lock-washer structure were preserved during the translocation burst, the motor would lose its register with the tracking strand after just one step, abolishing its strong grip for the substrate. On the other hand, if during the burst, the motor gradually converts from the lock-washer structure to a planar form as it translocates the substrate, it could partially preserve its grip on it throughout the process. Indeed, evidence that the motor can also adopt a planar structure has been recently found in x-ray and single-particle cryo-EM structures of the packaging ATPase of the closely-related phage asccphi28 obtained without DNA in the apo- and ADP-bound forms (Marc Morais, personal communication*). Based on these observations, we propose a translocation model in which, during its mechano-chemical cycle, the motor converts between a planar and a lock washer structure. A similar model has been proposed previously for this motor[3]. In this model, at the beginning of the dwell, the motor is in a planar structure that displays its weakest grip for the substrate because only the special subunit, numbered here as the first in the ring, is in direct contact with the tracking strand[4, 7, 20]. As exchange of ADP for ATP proceeds, hinge-like displacements between the first and second subunits, the second and third, and so on ensue, breaking the interface between the first and fifth subunits, resulting in a lock-washer structure (Fig. 3a). At the end of this process, each subunit now has established electrostatic interactions with phosphates in the tracking strand, which confer the motor its strong grip when fully loaded with ATP[4].These successive conformational changes are fixed and optimized to match the pitch of dsDNA. To explain how the motor matches the shorter pitch of the A-form helices, we propose that with these substrates, the displacement of the fifth subunit is partially interrupted by its earlier encounter with its corresponding phosphate in the tracking strand (Fig. 3b).

**Fig. 3.**
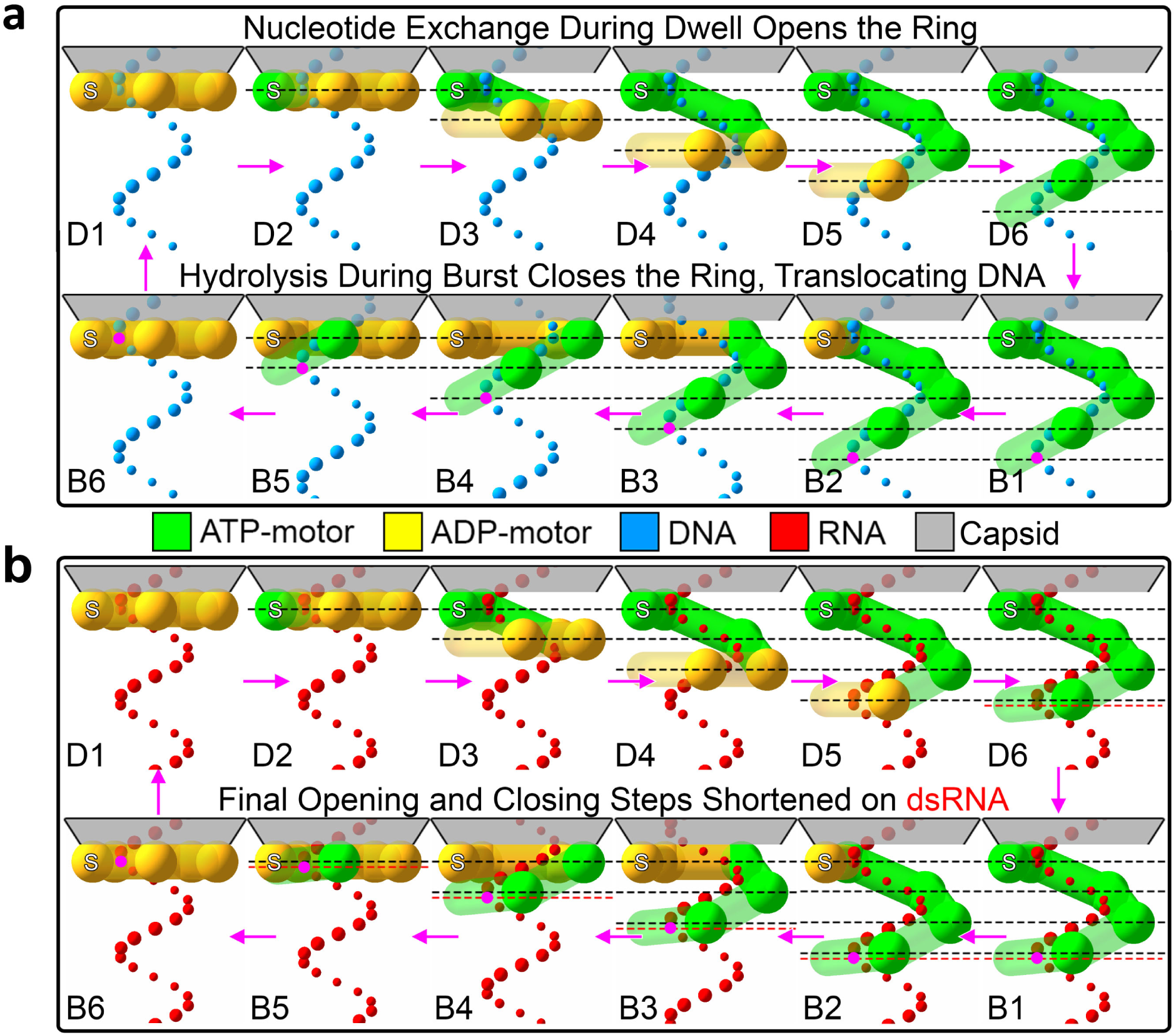
Translocation mechanism for the ϕ29 DNA packaging motor. The mechanism of the motor packaging (**a**) dsDNA and (**b**) dsRNA is shown. For each state, the ring ATPase complex is presented from the side, with each subunit represented as a cylinder with a sphere at its hinge and colored by nucleotide state. The capsid is above in grey and the special subunit is on the left, marked with an S. Tracking strand phosphates are depicted as colored spheres. The states in the dwell are labeled as D1-D6 and the states in the burst are labeled B1-B6. D6 and B1 are the same state, as well as D1 and B6, repeated only for horizontal comparison. Dashed lines with spacing 0.85 nm show the extent of ring opening. One phosphate is colored magenta during the burst to track translocation. During the dwell, ADP exchanges for ATP which opens the ring. During the burst, ATP hydrolysis drives DNA translocation. Note that the first exchange and hydrolysis is in the special subunit, which causes no ring opening or substrate translocation. On dsRNA, the mechanism is identical except for the final opening and closing steps, which are shorter and denoted by the red dashed lines. See also Supplementary Animation 1.

During the burst, each subunit, following hydrolysis and phosphate release[2], propels the substrate, sequentially returning to their conformation prior to ATP binding, progressively restoring the planar structure of the motor while translocating the substrate. The order in which the subunits hydrolyze their nucleotide during ring closing must be the same as that in which they exchanged nucleotide during ring opening, as this scheme preserves the greatest number of motor-substrate contacts (Fig. 3a), and is consistent with the highly coordinated mechano-chemical cycle previously proposed for this motor[5]. Each hydrolysis/phosphate-release event serves as the signal for the next subunit to hydrolyze its ATP, resulting in the four sequential steps that compose the burst. This model provides a straightforward explanation for the division of labor among the subunits, previously described for this motor. Specifically, the special subunit, proximal to the capsid and with a broken intersubunit interface, cannot undergo a power stroke; its ATP hydrolysis instead signals the beginning of the burst[5] (see Supplementary Discussion, Fig. 3a). By the time the distal (fifth) subunit executes its power stroke, the motor has regained its planar structure, and the phosphate in the tracking strand that was one full pitch down at the beginning of the cycle has been brought in register to contact the special subunit at the cycle’s end, most likely with the aid of some small substrate rotation[21]. With the dsDNA substrate, the burst is made up of four identical steps of 0.85 nm each (2.5 bp of dsDNA). With the A-form substrates, the last step of the burst is shorter due to the interrupted hinge movement of the fifth subunit during the dwell. The way the two ends of the open ring walk down the helical substrate is similar to the manner in which the two ends of an inchworm move as it walks – hence we name this model the *helical inchworm* (Fig. 3).

## Conclusion

Hand-over-hand mechanisms of translocation have been recently proposed for ring ATPase motors that adopt a lock-washer architecture and operate on disordered substrates. Here, we studied the dsDNA packaging motor of bacteriophage ϕ29 as a model of a ring-shaped translocase that operates on a substrate possessing a pre-existing structure. By using alternative double-helical structures we have revealed that the motor is capable of adapting itself to the periodicity of the substrate, and of maintaining the intersubunit coordination throughout its mechano-chemical cycle. This flexibility of the motor may reflect the need to accommodate the natural sequence-dependent structural variability of dsDNA[22]. We propose a model of translocation in which the ring converts between a lock-washer and a planar structure during its mechano-chemical cycle, in a way that maximizes its contacts with the spiral shape of the substrate. This model explains the high forces against which the motor can package the viral genome and its ability to resist the internal pressure that builds up in the capsid during packaging[21, 23, 24]. Interestingly, the ϕ29 motor has been shown to also package stretches of ssDNA[4]. It is possible that in this case the motor, while maintaining its translocation mechanism, imparts its helical symmetry on ssDNA, as has been shown for disordered substrate translocases. We speculate that the helical inchworm model proposed here may underlie the operation of other ring-shaped dsDNA translocases, as well as that of others that operate on disordered substrates.

## Materials and methods

### Modified substrates

Supplementary Fig. S5a shows a schematic of the substrate geometry. Two complementary ssRNA transcripts of 4 kb and 3.8 kb were generated using MEGAscript T7 vitro transcription (Themo-fisher Scientific) and purified using MEGAclear (Themo-fisher Scientific). The corresponding PCR templates were amplified from λ phage DNA (NEB), with ∼50% GC content; the same sequence is used for the 4 kb dsDNA substrate. For the DNA tracking strand hybrid, the 5’ end of the 4 kb transcript was biotin-labeled using Vaccinia Capping System (NEB) and 3-biotin-GTP (NEB) and purified using MEGAclear. The complementary DNA was synthetized using Protoscript II reverse transcriptase (NEB) and an adequate standard primer (IDT). For the RNA tracking strand hybrid the reverse transcription step is done using a 5’ biotinylated primer (IDT). For dsRNA the two transcripts were annealed at 65 °C for 15 min and cooled at a rate of 5 °C every 10 minutes until 25 °C, in 15 mM PIPES pH 6.5, 10 mM EDTA. The 187 nt overhang was completed using Protoscript II reverse transcriptase. For pulling experiments only the 5’-biotin-labeled 4 kb RNA transcript was used and for the corresponding reverse transcriptase reaction the primer was 5’-digoxigenin labeled.

### Optical tweezers packaging experiments

A dual trap optical tweezers instrument[3] was used to carry out single molecule packaging experiments using two polystyrene beads of 1 μm in diameter each. Trap stiffnesses were in the 0.4 – 0.7 pN/nm range. Raw data was acquired at 2.5 kHz.

ϕ29 prohead-motor complex[3, 4] was made by incubating purified proheads with recombinant motor ATPase (gp16) for 5 minutes at room temperature, after which ATPγS was added to a final concentration of 50 μM and kept for 24 hours at 4 °C.

Single packaging tethers were obtained following the *in situ* initiation protocol[3, 4] where one streptavidin-coated polystyrene beads (Banglabs) was previously deposited with substrate, while the second bead was deposited with prohead-motor-ATPγS complex. We used carboxyl-functionalized polystyrene beads (Banglabs) conjugated with antibody against gp8, a viral capsid protein.

*In situ* initiation of packaging occurs after few seconds of rubbing the two trapped beads in the presence of 0.25mM ATP. The packaging buffer (0.5x TMS) contained 25 mM Tris-HCl pH 7.8, 50 mM NaCl, 5 mM MgCl_2_ in 80% D_2_O, 20% H_2_O. [ATP] was 0.25 mM unless otherwise noted. 100 mg/ml glucose oxidase, 20 mg/ml catalase, and 5 mg/ml dextrose (Sigma-Aldrich) was used as oxygen scavenger system to increase tether lifetime.

Single molecule packaging trajectories were obtained by recording the decrease in tether extension (bead-to-bead distance) over time. Semi-passive mode was used to keep the force in a defined range, 7-12 pN (low force) or 30-35 pN (high force) unless otherwise notated. Force-feedback was used to characterize the step-wise slipping events in order to keep the force constant as the tether length increases.

Pulling experiments were performed over 4 kb fibers of the different polymers to characterize their mechanical properties[25-27] in the packaging buffer. Force vs extension pulling curves (Supplementary data Fig. S5b) were fitted to the extensible worm-like chain model, to experimentally obtain the persistence length (50 nm, 35 nm and 40 nm) and the stretch modulus (900 pN, 700 pN and 500 pN) for dsDNA, DNA/RNA hybrid and dsRNA, respectively. These parameters were used to convert bead extension to tether contour for packaging experiments.

### Data analysis

Raw packaging trajectories were converted to contour length using a method previously described[2, 3, 28]. Velocity distributions were obtained by binning position data filtered with a Savitsky-Golay differentiating filter of order 1, width 301 points (8.3Hz). These parameters were chosen because they smooth the dwell-burst cycle of the motor (∼11Hz) but retain enough definition to sharply capture the start of burst-sized slipping events.

Pairwise distributions were generated by taking the autocorrelation of traces filtered by a moving average filter of 41 points (61Hz). Distributions were graded by their periodicity via a method previously described[3], and the top 30% of distributions were selected for averaging.

Burst-sized slipping was analyzed by using the Kalafut-Visscher stepfinding algorithm[29] to find regions of increasing tether length. High-force stepfinding was done using a hidden markov model-based stepfinding algorithm[30].

## Supporting information

Supplementary Animation 1

## Data Availability

The data that support the findings of this study are available from the corresponding author upon reasonable request.

## Code Availability

The code that was used in this study is available at https://doi.org/10.5281/zenodo.3718873 under the GNU GPLv3 license, and Alexander Tong may be contacted for details upon reasonable request.

## Acknowledgments

We thank all members of the Bustamante laboratory for insightful discussions. This research was supported by the Nanomachines program (KC1203) funded by the Office of Basic Energy Sciences of the U.S. Department of Energy (DOE) contract no DE-AC02-05CH11231 and by the National Institute of Health grant R01GM032543; C.B. is a Howard Hughes Medical Institute investigator. J.P.C. acknowledges support from the PEW Latin American Fellows program in the biomedical sciences.

## Author contributions

J.P.C. and C.B. conceived the project; J.P.C. designed and prepared substrate constructs; J.P.C., A.T. and S.T. performed single molecule packaging experiments; A.T. analyzed the data; P.J. provided critical reagents for the experiments; C.B., J.P.C. and A.T. wrote the paper.

* The lock-washer structure found for the ϕ29 packaging motor by the group of Marc Morais is available as a preprint in bioRxiv, submitted in parallel with this preprint.

