## Supplementary Material for "A DNA translocase operates by cycling between planar and lock-washer structures"

#### Supplementary discussion

##### The mechano-chemistry of the motor is preserved with A-form substrates

It has been described that in the dwell-burst cycle of the packaging motor the dwell times $\tau$ follow a gamma distribution, indicative of multiple rate-limiting events. These events were identified as ADP release by each subunit, which occurs sequentially around the ring[1, 2]. The dwell time distributions observed with the A-form substrates are also gamma-distributed with similar scale and shape parameters (Supplementary Fig. S1). However for the RTS hybrid and dsRNA, the apparent number of rate limiting event, using the estimator $n_{min} = \langle\tau\rangle^2/(\langle\tau^2\rangle - \langle\tau\rangle^2)$, is 4.67 and 4.68 respectively, marginally smaller than for dsDNA and the DTS hybrid, which are 5.05 and 5.04, respectively (Supplementary Fig. S1). Such a small change in $n_{min}$ may reflect a slight decrease in the ADP release rate by one of the subunits, for example, that of the special subunit because of its modified interaction with the RNA tracking strand. These differences notwithstanding, the mechano-chemical scheme of the motor is preserved when packaging alternative substrates.

##### Differing resolution limits with substrates

In dual-trap optical tweezers, the signal-to-noise ratio (SNR) is determined by the size of the beads used in the experiment and the stiffness *k* of the tether held between them[3]. The size of the bead cannot go appreciably below 1 µm, the size used in these experiments, and therefore the resolution is proportional to *k*. Under high forces (>10 pN), the substrates are fully extended and *k* is given by the stretch modulus of the tether (pN) divided by its length (nm). The measured stretch moduli of the dsDNA, hybrid, and dsRNA polymers are 900, 700, and 500 pN, respectively (Supplementary Fig. S5b). Therefore, the SNR of the hybrid and dsRNA is 0.78 and 0.56 times that of dsDNA, which explains why the data on the alternative substrates is noiser than that of dsDNA, and why it makes the resolving of the small 0.45 nm (0.15 nm) correction step for the hybrid (dsRNA) more difficult.

##### Analysis of reverse translocation events

We propose that during the reverse translocation events the motor transitions into a packaging-incompetent (PI) state that has reduced grip for the substrate. From this state, the motor can either return to the packaging-competent (PC) state (in which case, the event is indistinguishable from a pause) or proceed to slip. Each reverse translocation event is divided into three sections: the *start dwell*, where the motor transitions into the PI state before burst-sized slipping starts, the *slipping dwells*, which separate the burst-sized increases in tether length, and the *end dwell*, where the burst-sized slipping ends and the motor transitions back to the PC state (Supplementary Fig. S2a). To investigate the process by which the motor transitions from the PC state to the PI state, we analyzed the effect of high (30 pN) and low (10 pN) opposing force, as well as the effect of high (250 μM) and low (25 and 10 μM) [ATP] on the dynamics of this process. Only events that recovered packaging after slipping were analyzed; those that exhibited large slipping were rejected, as they are indistinguishable from normal, non-burst-sized slipping.

Limiting [ATP] increases the frequency of reverse translocation events, indicating that the transition from the PC to the PI state takes place when the motor is not fully saturated with ATP (Supplementary Fig. S2b). On the other hand, the duration of the start dwell and the total extension of the reverse translocation event are independent of [ATP], implying that while in the PI state, the motor is unable to exchange ADP for ATP (Supplementary Fig. S2b). Moreover, these observations suggest that during a regular packaging dwell, a kinetic competition ensues between the completion of ATP exchange in the PC state and the attainment of the PI state (Supplementary Fig. S3).

Because the slipping dwells are much shorter than the start dwell, we conclude that the motor is in a different state during slipping than when it first entered the PI state during the start dwell. Therefore, at some point during the start dwell, the motor must make a transition from a “PI-gripping” state to a “PI-slipping” state (Supplementary Fig. S3). Consistent with this interpretation, we find that the duration of the start dwell is gamma-distributed, supporting the existence of transitions among several states during this time (Supplementary Fig. S2c). Incidentally, on the DTS hybrid we observe paused states (Fig. 1d and 2a, Supplementary Fig. S4), but do not observe reverse translocation events, indicative that, with this substrate, the motor can enter the PI-gripping state but not the PI-slipping state. This observation suggests that the transition from the PI-gripping to the PI-slipping state results from a sub-optimal interaction between the motor and the RNA tracking strand. Furthermore, the fact that opposing force decreases the start dwell duration and increases the frequency of reverse translocation events (Supplementary Fig. S2b), indicates that force favors the transition from the PC to the PI-gripping state, and/or from the PI-gripping to the PI-slipping state (Supplementary Fig. S3b). Analysis of the distribution of slipping dwells reveals that at low force the dwells are gamma distributed, while at high force they follow a single-exponential distribution, indicating that slipping involves at least two rate limiting steps, one more force-dependent than the other (Supplementary Fig. S2c).

The fact that the total extension of reverse translocation events does not change with opposing force (Supplementary Fig. S2c), whereas the slipping dwells become shorter, implies that the transition out of the PI-slipping state occurs when the motor reattaches to the substrate (Supplementary Fig. S3). Were this not the case, and the transition would have happened during the end dwell, the total extension of the reverse translocation event would have increased with opposing force. Additionally, the length of the end dwell is longer than a regular dwell, indicating that during this dwell, the motor first enters a state that can stably grab onto the substrate but cannot translocate (Supplementary Fig. S2b).

Finally, we examined how the motor transits back to PC state during the end dwell. The end dwell is lengthened at limiting [ATP] (Supplementary Fig. S2b), indicating that the motor, not fully saturated with ATP, transits from the PI-gripping state to the PC state where it can restart its halted nucleotide exchange and resume packaging (Supplementary Fig. S3). We attribute the extra time spent in the PI-gripping state before packaging resumes to a loss of communication in the ring: under normal circumstances, there is a signaling cascade that accompanies nucleotide exchange[4], whereby binding of substrate to the special subunit causes the first ATP to be exchanged, which sends a signal to next subunit to exchange its ATP, and so on until the ring is saturated. However, at the beginning of the end dwell, the motor has already some ATP bound and although the special subunit has just re-engaged with the substrate, the signaling from this subunit can no longer effectively resume the nucleotide exchange cascade. The completion of the slow nucleotide exchange event restores the signaling cascade and transitions the motor from the PI-gripping state to the PC state.

Additionally, we observe pauses that do not result in burst-sized slipping. They have a duration of 250 ± 5 ms (median ± s.e.m.), which is close to the sum of the lengths of a start and an end dwell, minus a slipping dwell. This observation confirms that such pauses occur when the motor transitions from the PC to the PI-gripping state and back without visiting the PI-slipping state. The frequencies of pauses and reverse translocation events increase equally with low [ATP] (Supplementary Fig. S4b), which supports the idea that to enter a pause, the motor must also transition to the PI-gripping state. Moreover, the fact that the ratios of these frequencies are independent of [ATP] indicates that the transition from the PI-gripping state to the PI-slipping state must be nucleotide-exchange independent, which is, in turn, consistent with the proposition that ATP exchange is halted when the motor is in any of its PI states (slipping or gripping).

Taken together, all the above observations lead us to conclude that in the PI state the motor-substrate interaction has been compromised resulting in the lack of the signals needed for nucleotide exchange and hydrolysis cascade[4]. While in the PI-slipping state, the motor, in its ATP-unsaturated state, is unable to hold onto the RNA tracking strand, slipping a full turn of the helix successively. The inability of the motor to make the proper phosphate contacts with the tracking strand of the A-form substrates explains the interruption of the mechano-chemical events of the motor's cycle.

**ATP hydrolysis cascade initiation by the special subunit**

The strict order in which the mechano-chemical processes occur in this motor is paramount to its coordination, which in the helical inchworm model proposed here can be accomplished through trans-acting residues for all steps except for the initiation of the ATP hydrolysis cascade after the last subunit binds ATP[4]. For this step, a signal must be sent somehow from the subunit distal to the capsid, across the broken interface, to the special subunit. We attribute this to stress being built up in the motor subunits as the motor opens along the helix. By the time the last subunit has bound ATP, this stress buildup is sufficient to trigger hydrolysis by the special subunit, thus initiating the translocation cascade. This model also explains the spontaneous triggering of the translocation burst observed under very low [ATP], where the motor has not yet fully exchanged ADP for ATP[4]. Presumably, the sub-critical stress buildup in the ring at partial saturation allows the special subunit to eventually overcome the activation barrier required for firing when the motor has at least 3 subunits bound with ATP, since we never observe spontaneous triggering that results in a single translocation step, which would occur if 2 subunits were bound to ATP.

**Supplementary Figures**

### Fig. S1

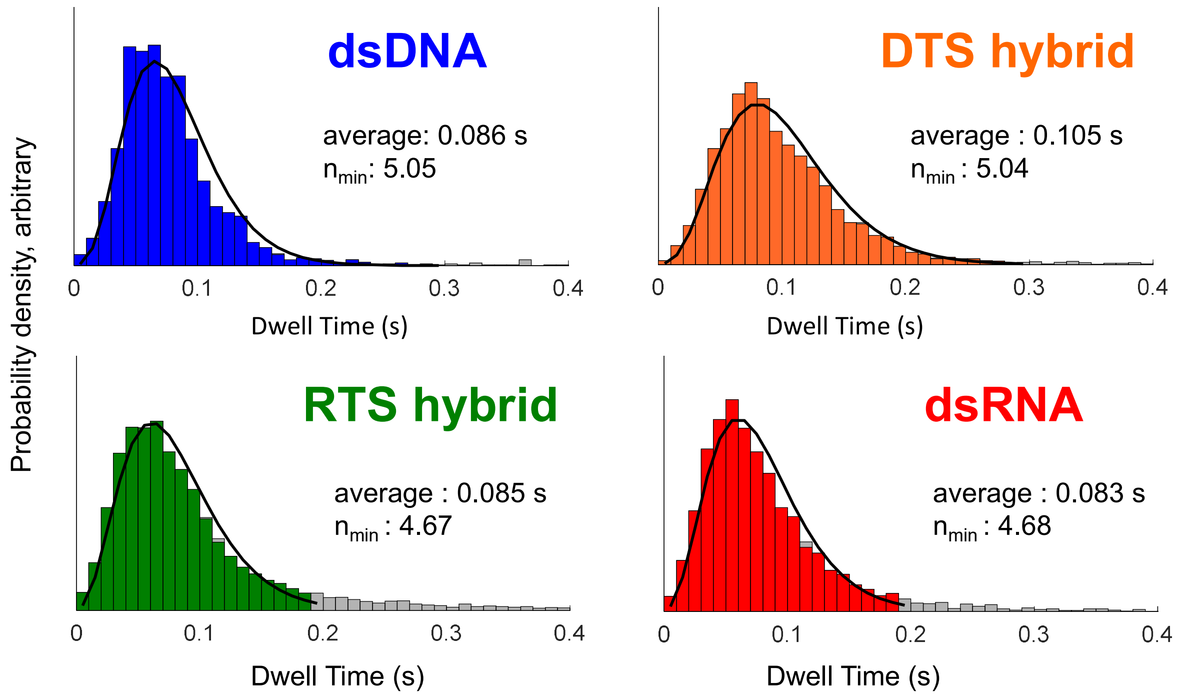


**Supplementary Fig. S1|Dwell time distributions.** The dwell times of the four substrates are obtained from stepfinding and are fit to a gamma distribution. The mean and the apparent number of rate limiting steps $N_{min} = \langle\tau\rangle^2/(\langle\tau^2\rangle - \langle\tau\rangle^2)$ are noted.

Fig. S2

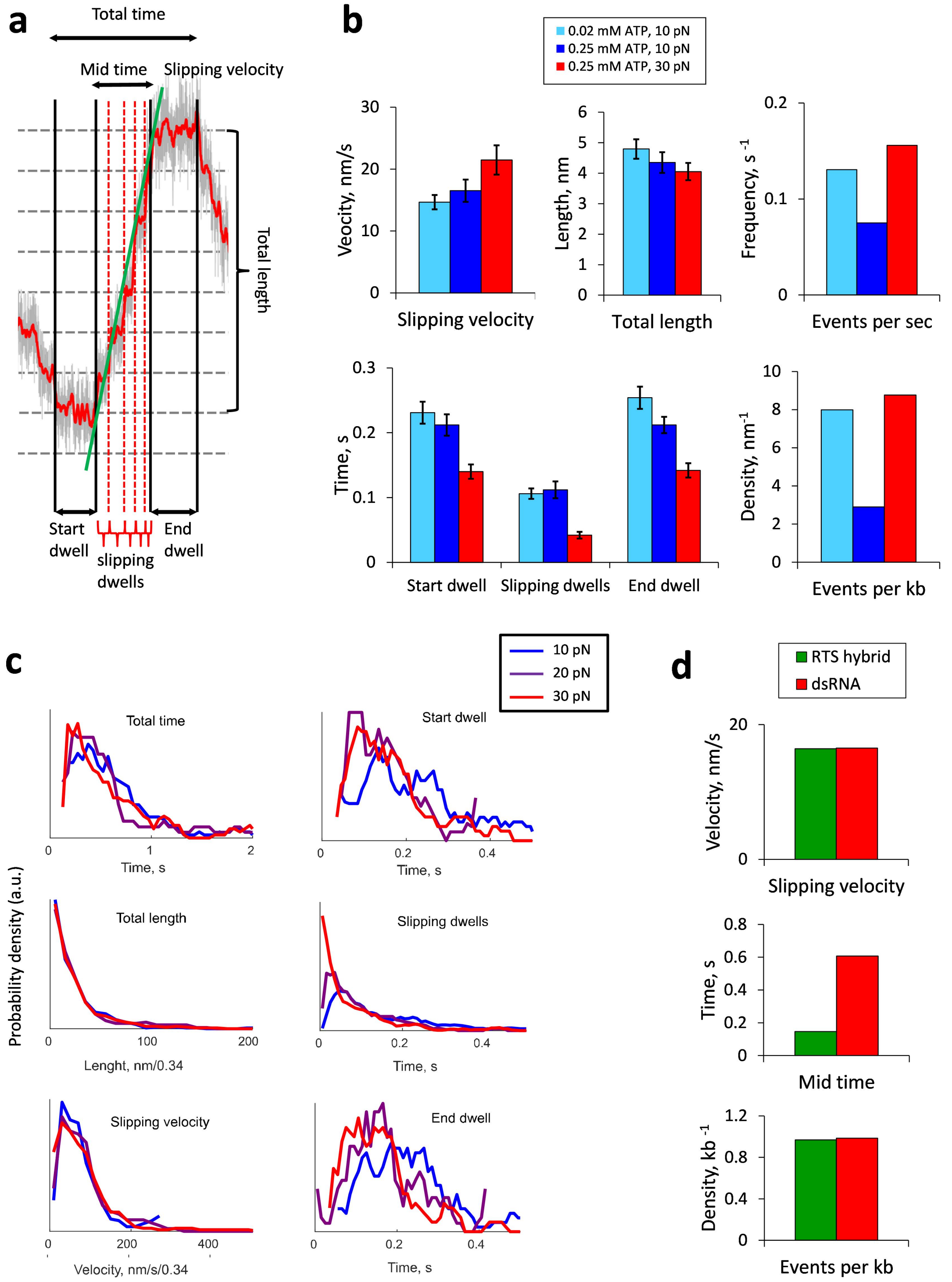

a
Total time
Mid time
Slipping velocity
Total length
Start dwell
slipping dwells
End dwell
b
0.02 mM ATP, 10 pN
0.25 mM ATP, 10 pN
0.25 mM ATP, 30 pN
Vecocity, nm/s
Slipping velocity
Length, nm
Total length
Frequency, s -1
Events per sec
Time, s
Start dwell
Slipping dwells
End dwell
Density, nm-1
Events per kb
c
10 pN
20 pN
30 pN
Probability density (a.u.)
Total time
Time, s
Start dwell
Time, s
Total length
Lenght, nm/0.34
Slipping dwells
Time, s
Slipping velocity
Velocity, nm/s/0.34
End dwell
Time, s
d
RTS hybrid
dsRNA
Velocity, nm/s
Slipping velocity
Time, s
Mid time
Density, kb -1
Events per kb

**Supplementary Fig. S2|Reverse translocation dynamics. a**. An individual event is divided into three sections to differentiate the first and last dwell from the dwells during burst-sized slipping (slipping dwells). **b**. Medians of select aspects of the reverse translocation events on dsRNA are shown, error bars are s.e.m.. **c**. Distributions of various aspects of the events are shown under varying opposing forces. **d**. Differences of select statistics are shown when the substrate is the RTS hybrid compared to dsRNA. Unless otherwise noted, [ATP] is 0.25 mM and the force is 10 pN.

### Fig. S3

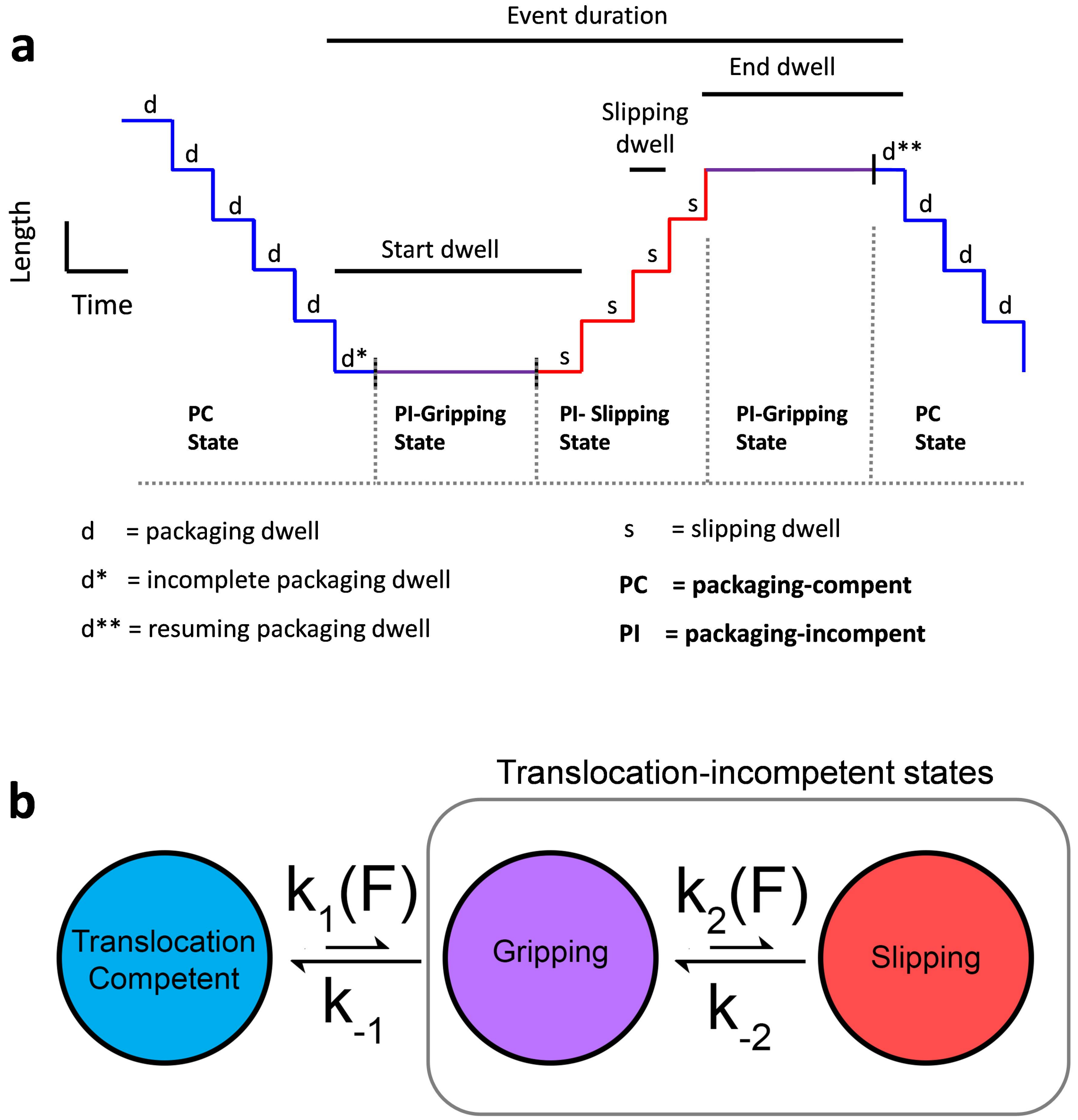


**Supplementary Fig. S3 | Model of the reverse translocation events. a**. A schematic packaging trajectory of a reverse translocation event is shown, with underlying states labeled. Blue, purple, and red lines correspond to the motor being in the packaging-competent (PC), packaging-incompetent (PI)-gripping, and PI-slipping states, respectively. **b**. The state diagram of the motor is depicted, showing rates and their relative sizes and force dependence. Note that the transition to the gripping and slipping states is force-dependent ($k_1(F)$, $k_2(F)$), and from the gripping state the motor is more likely to transition out of the translocation-incompetent state (leading to a pause) than proceed to slip ($k_2 < k_{-1}$).

### Fig. S4

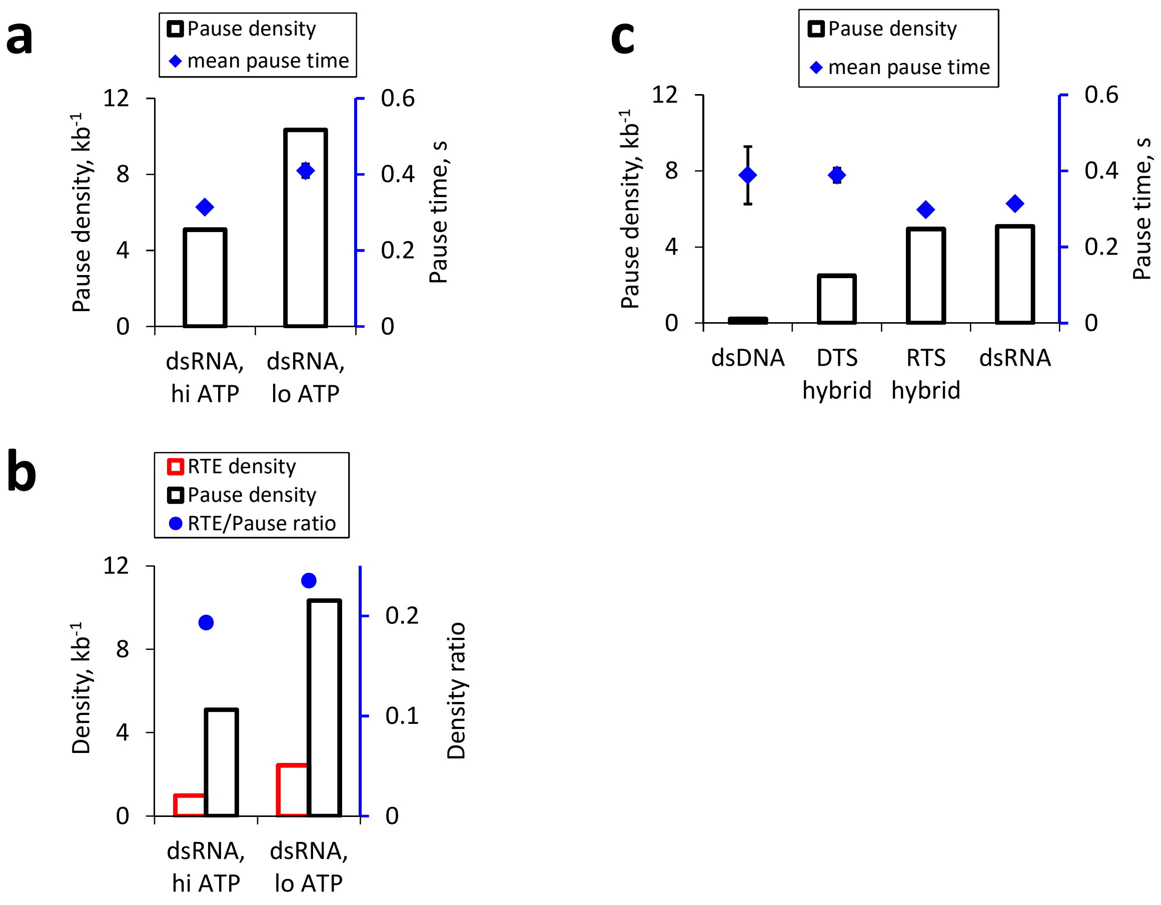


**Supplementary Fig. S4 | Motor pausing dynamics. a**. Limiting [ATP] increases pause density appreciably but only slightly increases pause time. **b**. Limiting [ATP] increases proportionally both pause and reverse translocation event (RTE) density on dsRNA. **c**. The effect of substrate identity on pause time and density is shown. For the substrates that show appreciable amounts of pausing (the A-form ones), their pause times are similar. For all plots, [ATP] = 0.25mM unless otherwise noted, F = 10 pN; bars refer to the left axis and dots refer to the right axis.

### Fig. S5

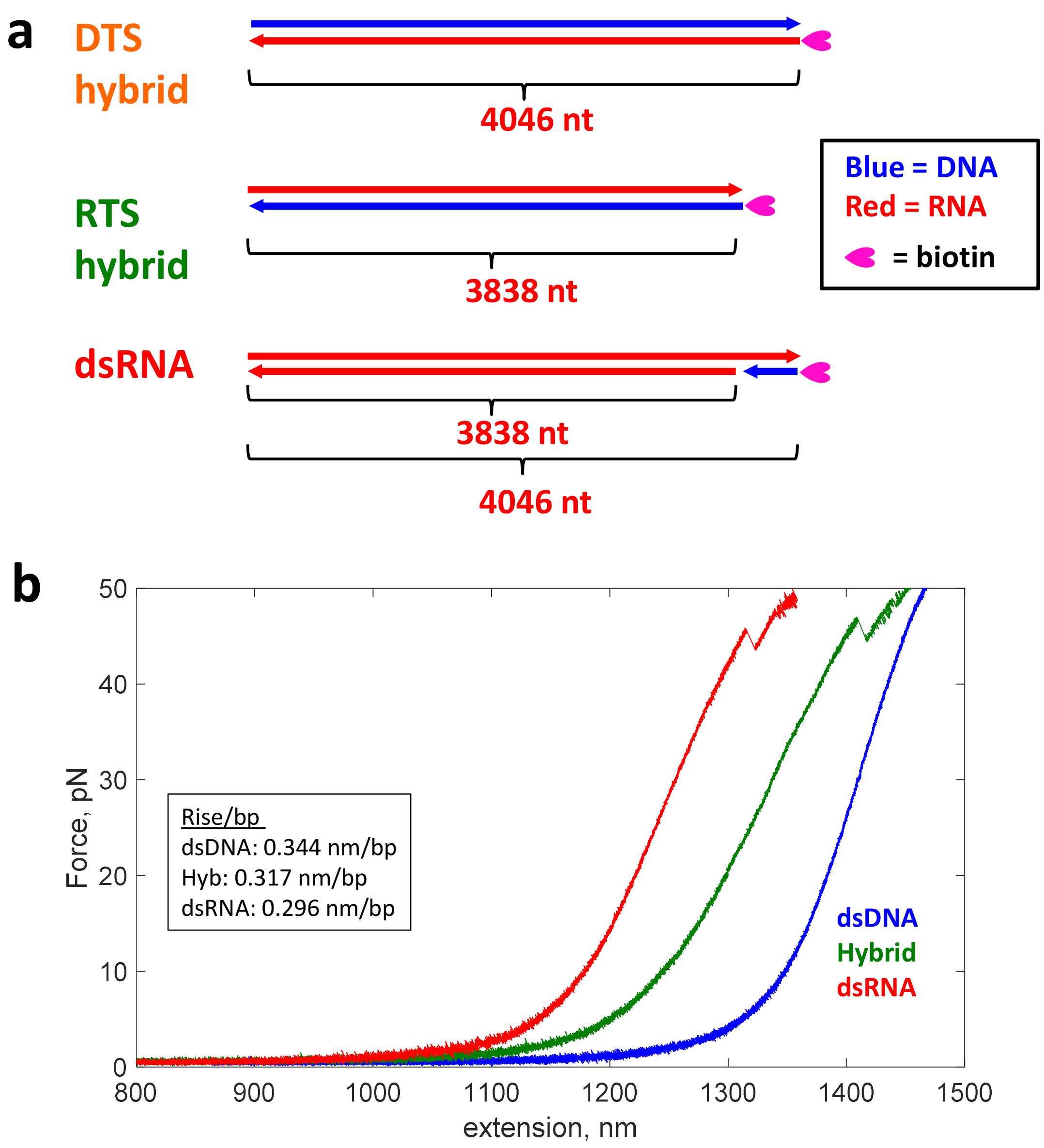


**Supplementary Fig. S5 |Alternative A-form substrates construction and characterization. a.** The construction of the alternative substrates is shown, where red solid lines represent RNA strands and blue lines represent DNA strands synthetized using reverse transcription (see methods). Arrow heads indicate 5’-to-3’ direction. **b**. Force vs. Extension plot obtained from pulling experiments performed on 4 kb constructs of the different substrate polymers. DNA/RNA and dsRNA have a diminished contour length (left shift in the curves) compared to dsDNA, which means a smaller rise per base pair (see inset).

**Supplementary animation 1.** Helical inchworm mechanism of translocation. For legend refer to figure 3.
