## Supplementary figures and images for "A DNA translocase operates by cycling between planar and lock-washer structures"

### Supplementary Animation 1

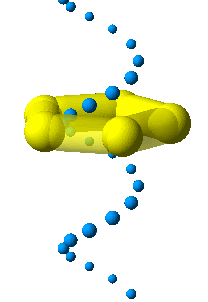
